# E3 Ligase UBR5 HECT domain mutations in lymphoma control maturation of B cells via alternative splicing

**DOI:** 10.1101/732180

**Authors:** Samantha A. Swenson, Tyler J. Gilbreath, Heather Vahle, R. Willow Hynes-Smith, Jared H. Graham, Henry Chun Hin Law, Nicholas T. Woods, Michael R. Green, Shannon M. Buckley

## Abstract

Coordination of a number of molecular mechanisms including transcription, alternative splicing, and class switch recombination are required to facilitate development, activation, and survival of B cells. Disruption of these pathways can result in malignant transformation. Recently, next generation sequencing has identified a number of novel mutations in mantle cell lymphoma (MCL) patients including the ubiquitin E3 ligase UBR5. Approximately 18% of MCL patients were found to have mutations in UBR5 with the majority of mutations within the HECT domain of the protein which can accept and transfer ubiquitin molecules to the substrate. Determining if UBR5 controls the maturation of B cells is important to fully understand malignant transformation to MCL. To elucidate the role of UBR5 in B cell maturation and activation we generated a conditional mutant disrupting UBR5’s C-terminal HECT domain. Loss of the UBR5 HECT domain leads to a block in maturation of B cells in the spleen and up-regulation of proteins associated with mRNA splicing via the spliceosome. Our studies reveal a novel role of UBR5 in B cell maturation by regulating alternative splicing of key transcripts during B cell development and suggests UBR5 mutations may promote mantle cell lymphoma initiation.

**KEY POINTS:**

- Utilizing a novel mouse model mimicking MCL patient mutations, the loss of UBR5’s HECT domain causes alterations in B cell development.
- UBR5 mutations lead to stabilization of UBR5 and aberrant splicing.

## INTRODUCTION

Mantle Cell Lymphoma (MCL) is a rare, aggressive form of Non-Hodgkin’s Lymphoma (NHL).^1^ Although MCL represents only ~6% of NHL lymphoma cases, it has one of the highest mortality rates of all lymphomas with only a 50% five year survival.^2^ Given the high mortality rate and propensity for recurrence, having better comprehension of mutations found in MCL and how disease develops in B cells will open avenues for identifying new therapies. Recently, the ubiquitin protein ligase E3 component n-recognin 5 (UBR5) was found mutated in ~18% of patients with MCL.^3^ The majority of mutations identified in *UBR5* were frame shift mutations found within its HECT domain, which can accept and transfer ubiquitin molecules to the substrate, leading to a premature stop codon prior to the cysteine residue associated with ubiquitin transfer.

UBR5 is a large ~300kDa protein HECT E3 ligase with a conserved carboxyl-terminal HECT domain. In HECT E3 ligases, the N-terminal portion (N-lobe) of the enzyme interacts with E2 ubiquitin-conjugating enzymes and determines substrate specificity while the C-terminal HECT domain (C-lobe) contains a catalytic cysteine residue that binds ubiquitin.^4^ The two lobes are connected by a flexible linker that allows for shifting orientation between N- and C-lobes during ubiquitin transfer to allow for efficient movement of ubiquitin from the E3 ligase to the substrate protein. UBR5 regulates a number of cellular processes including metabolism, apoptosis, angiogenesis, gene expression, and genome integrity.^5–10^ Overexpression of UBR5 has been found in a number of cancers including ovarian, breast, hepatocellular, squamous cell carcinoma, and melanoma.^11–14^

Determining if, and at what stage – transcriptional, translational, or proteomic – UBR5 controls maturation of B cells is important for fully understanding B cell development and lymphoma transformation. In order to elucidate the role of UBR5 in B cell maturation and activation, we generated a conditional mutant disrupting the C-terminal HECT domain. Loss of the HECT domain leads to a block in maturation of B cells, and follicular B cells are phenotypically abnormal with low expression of IgD and high expression of IgM. Upon immune stimulation, B cells lacking the HECT domain show decreased germinal center formation and reduced antibody producing plasma cells suggesting both defects in phenotype and function of mature B cells. Proteomic studies reveal up-regulation of proteins associated with mRNA splicing via the spliceosome and indicates that UBR5 interacts with splicing factors (SF3B3, SMC2, PRPF8, DHX15, SNRNP200, and EFTUD2). Our studies reveal a novel role of UBR5 in B cell maturation by regulating alternative splicing of key transcripts during B cell development and suggests *UBR5* mutations in MCL lead to disease initiation.

## METHODS

### Mice

*Ubr5*HECT mutant mice were developed using Easi-CRISPR as previously published^15^ and crossed with *Mb1*^*CRE*^ mice (kind gift of Michael Reth). Floxed *Ubr5* alleles were validated by PCR using primers designed to amplify targeted alleles (supplemental materials and methods). For immune stimulation, mice were immunized by intraperitoneal injection with 1×10^8^ sheep’s red blood cells (SRBC) (Innovative Research). All mice were housed in a pathogen-free facility at University of Nebraska Medical Center. Procedures performed were approved by Institutional Animal Care and Use Committee of University of Nebraska Medical Center in accordance with NIH guidelines.

### B cell isolation and culture

Total bone marrow (BM) and splenocytes were isolated from 6-week-old mice. B220^+^ cells were isolated using MojoSort Streptavidin Nanobeads (Biolegend, San Diego, CA) following manufacturer’s protocol. B cells were cultured in RPMI 1640 Base Media (Hyclone), 10% fetal bovine serum (Atlanta Biologicals), 2mM L-glutamine (Corning), 50µM 2-mercaptoethanol (Corning), 20mM HEPES (4-(2-hydroxyethyl)-1-piperazineethanesulfonic acid) (Hyclone), and 1X penicillin/streptomycin. Cycloheximide (Millipore) was used at a final concentration of 10µg/mL.

### FlowCytometry analysis

For flowcytometry analysis cells were stained for 1 hour in 3% FBS in PBS. Antibodies found in supplemental materials and methods. For cell cycle analysis, cells were fixed and permeabilized following Biolegend intracellular staining protocol and stained with Ki67 and DAPI (4’,6-diamidino-2-phenylindole).

### ELISA

Mice were bled on Day 0 and Day 8 following immune stimulation. Serum was collected following Abcam ELISA sample preparation guide, diluted 1:10 with PBS. SRBC, diluted 1:10 in PBS, was used as a control for cross reactivity of antibodies due to presence of SRBC. ELISA was performed with positive reference antigen mixture, and PBS as a negative control according to manufactures directions (BD Pharmingen).

### Histological Staining

Mouse spleens were fixed in 10% (vol/vol) buffered formalin phosphate for 24 hours then placed in 70% ethanol. Sections were stained with H&E using standard protocols. Spleen sections were stained in a 1:500 dilution of UBR5 antibody ab70311 (Abcam) and Ki67. For germinal center analysis, mice were immunized with SRBC and sacrificed eight days later. Spleen sections were stained in a 1:500 dilution of biotinylated peanut agglutinin (PNA) antibody B-1075 (Vector Labs). Images were captured with a Zeiss Observer.Z1 microscope (Zeiss International, Germany) and Zen Pro software (Zeiss International, Germany) was used to process the images.

### Nuclear Fractionation

Nuclei from 293Ts (ATCC) were collected as described previously.^16^ Nuclei were lysed in low salt buffer (10mM Tris-HCl pH7.4, 0.2mM MgCl_2_, 1% Triton-X 100) containing protease and phosphatase inhibitors for 15 minutes on a rocker at 4ºC. 10-30% glycerol gradients were prepared as previously described.^17^ Samples were spun at 28,000rpm for 13 hours at 4ºC. 200uL fractions were collected for a total of 25 fractions.

### Western Blot and Immunoprecipitation Analysis

For western blot analysis, samples were lysed in Pierce RIPA buffer containing protease and phosphatase inhibitor cocktail (ThermoFisher). Samples were separated by SDS-PAGE, transferred onto PVDF (Millipore), and blocked in 5% milk. Antibodies were prepared in 5% BSA. Horse Radish Peroxidase conjugated secondary antibodies (Jackson laboratories) were prepared in 5% milk.

### RNA extraction and quantitative real-time PCR

Total RNA was harvested using QIAGEN RNeasy Kit (QIAGEN, Hilden, Germany). Following RNA extraction, cDNA synthesis was performed using High Capacity RNA-to-cDNA Kit (ThermoFisher). qRT-PCR was carried out on equal concentrations of cDNA for each sample using iTaq Universal SYBR Green Supermix. All primers can be found in supplemental materials and methods.

### Mass Spectrometry

For global proteome quantification, B220^+^ splenocytes were isolated from 3 mice per genotype. Samples were prepared for and TMT labeled per manufacturer’s protocol (ThermoScientific TMT10plex Mass Tag Labeling Kits). Following TMT labeling, acetonitrile was removed by speedvac and samples were resuspended in 0.1% trifluoroacetic acid (TFA). Sample cleanup with C18 tips was performed per manufacturer’s protocol (Pierce). Sample concentrations were re-quantified (Pierce Quantitative Colorimetric Peptide Assay kit) and then combined in equal concentration. Following combination, samples were dried by speedvac and fractionated by ThermoScientific high pH reverse phase fractionation kit following manufacturer’s protocol for TMT. Resulting fractions were run in a speedvac to dryness and resuspended in 0.1% Formic Acid for mass spectrometry (MS) analysis (see supplemental methods). Data are available via ProteomeXchange with identifier PXD014307.

For immunoprecipitation, cells were lysed in 20mM Tris pH 7.5, 150mM NaCl, 1mM EDTA, and protease and phosphatase inhibitors. Immunoprecipitation was performed overnight at 4°C using anti-UBR5 antibody from Cell Signaling (8755) or rabbit IgG control. Protein A agarose beads (Cell signaling) were used for IgG pulldown. Samples were washed 5X with lysis buffer prior to analysis by MS.

### Statistical analysis

All experiments were performed in triplicate unless noted and statistical analyses were performed using paired two-tailed Student’s t-test assuming experimental samples of equal variance. * P<0.05, ** P<0.01, *** P<0.001, **** P<0.0001

## RESULTS

### *Ubr5* mutations are specific to MCL

Meissner, B. et al originally identified UBR5 mutations in ~18% of MCL patients.^3^ In recent cross-sectional genomic profiling of multiple lymphoma subtypes, we identified UBR5 as one of 8 genes that had a significantly higher frequency of mutation in MCL compared to other lymphoma subtypes. These mutations were observed in 21 out of 196 MCL tumors (10.7%) and 15 out of 559 tumors (2.7%) of other histologic subtypes. However, mutations within the HECT domain were found only in MCL tumors (Figure 1B).^18^ Mutations within the HECT domain of UBR5 are therefore a disease-specific genetic feature of MCL.

**Figure 1.**
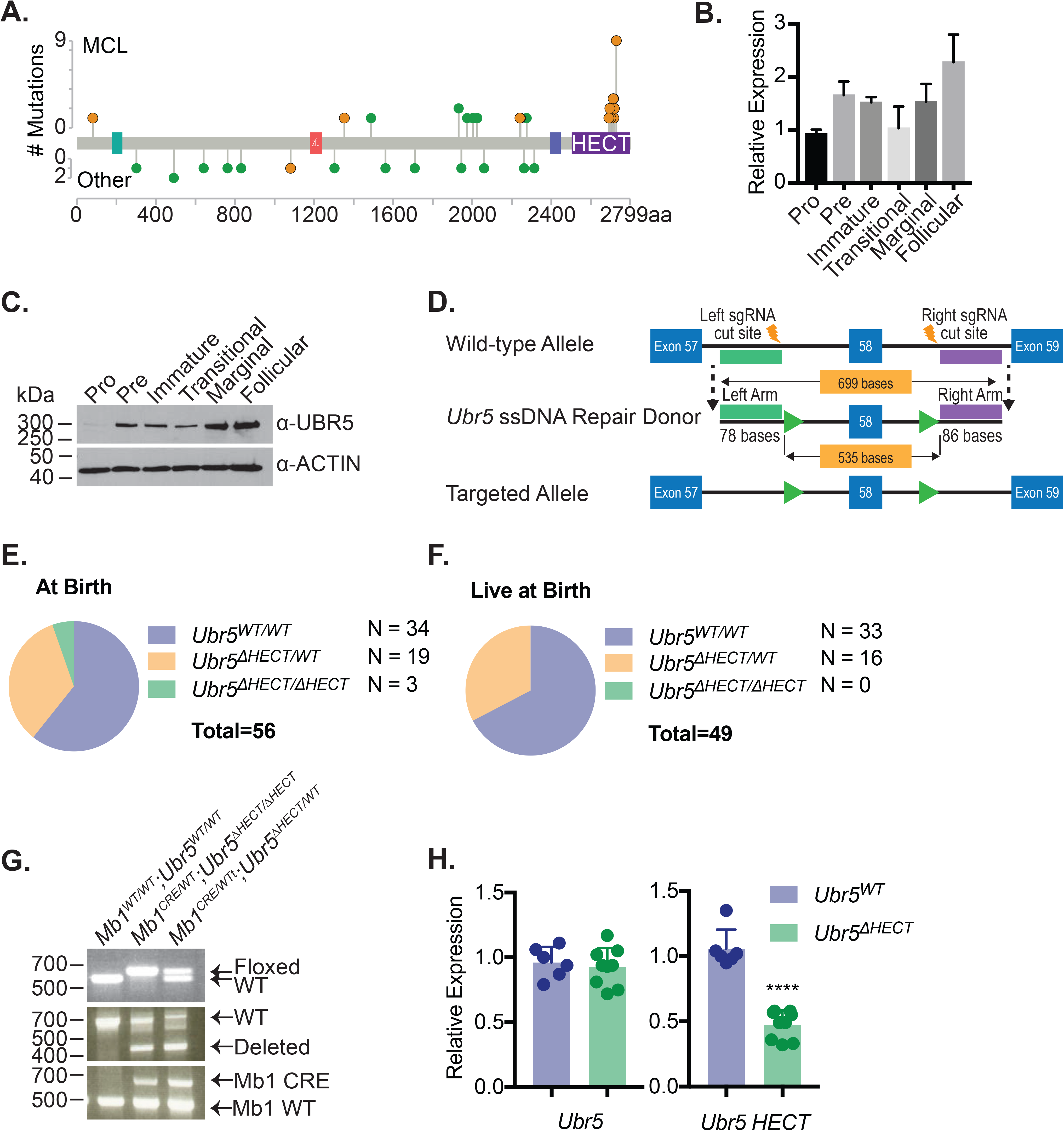
Expression of *Ubr5* in B cells and generation of conditional *Ubr5* HECT domain knockout model. (A) Frequency of UBR5 mutations in lymphoma patients. orange = nonsense/frameshift; green = missense. (B) Relative qRT-PCR and (C) Western blot expression of UBR5 within different B cell populations: pro B cells (B220^+^IgM^−^ckit^+^), pre B cells (B220^+^IgM^−^CD25^+^), immature B cells (B220^+^IgM^lo^IgD^−^) from the BM, and transitional (B220^+^CD93^+^), marginal zone B cells (B220^+^CD21^+^CD23^−^), and follicular B cells (B220^+^CD21^+^CD23^+^) isolated from the spleen of 6 weeks old WT C57Bl6 mice. (D) Schematic of targeting strategy used to insert loxP sites flanking exon 58 of *Ubr5*. (E) Distribution of pups born from crossing *Ubr5*^*ΔHECT/ΔHECT*^ with E2A^CRE^ (F) Pups alive at birth per genotype. (G) Genotyping PCR of targeted alleles and deletion in six-week old mice bred to B lymphocyte specific *Mb1*^*CRE*^ mouse model. PCR was performed on BM. (H) Relative expression of *Ubr5* and *Ubr5* HECT domain by qRT-PCR in spleen B220^+^ cells.

### Generation of a conditional UBR5 HECT domain mutant

Since the role of *Ubr5* in lymphopoiesis is unknown, we first evaluated the expression of *Ubr5* in B cell sub-groups during development by purifying pro, pre, and immature B cells from BM of 6-week-old C57/BL6 wild-type (WT) mice. Additionally, transitional, follicular, and marginal zone B cell populations were purified from spleens. The pro B cell population showed lowest expression of *Ubr5* at both the RNA and protein level, whereas the highest expression of *Ubr5* was found in mature splenic populations (follicular and marginal B cells) (Figure 1B-C). These studies suggest a role for UBR5 at different stages of B cell development.

Saunders *et al.* generated a *Ubr5*-null mouse; unfortunately, these mice die during embryogenesis prior to onset of definitive hematopoiesis at E10.5.^19^ In order to understand the role of UBR5 mutations in B cell development, we generated a conditional allele targeting exon 58 that would lead to a truncated protein lacking a critical cysteine required for ubiquitin conjugation in the HECT domain and mimicking mutations found in MCL patients (Figure 1D).^20^ We first crossed our *Ubr5*^*ΔHECT/+*^ mice to *E2A^CRE^*;*Ubr5*^*ΔHECT/+*^ mice that delete in early embryogenesis.^19, 21^ As with null mice, *E2A*^*CRE*^;*Ubr5*^*ΔHECT/ΔHECT*^ were not viable. In contrast to *Ubr5*-null mice, 3 of 48 pups found dead at birth were *E2A*^*CRE*^;*Ubr5*^*ΔHECT/ΔHECT*^, suggesting the HECT domain is not required for embryogenesis and early stages of hematopoiesis (Figure 1E-F).

Since *E2A*^*CRE*^;*Ubr5*^*ΔHECT/ΔHECT*^ mice die prior to or at birth, to study B cell development we crossed *Ubr5*^*ΔHECT/∆HECT*^ mice to *Mb1*^*WT/CRE;*^ *Ubr5*^*ΔHECT/∆HECT*^, which deletes in early B-lymphocytes (Figure 1G).^22^ To determine the specific loss of the HECT domain, we performed qRT-PCR with primers flanking the HECT domain and primers amplifying a region on the N-terminus in B220^+^ splenocytes. qRT-PCR showed no decrease in *Ubr5* expression, but significant decrease in *Ubr5* HECT domain expression (Figure 1H). These studies demonstrate the loss of HECT domain of UBR5 targeted mice.

### Impaired B cell maturation following deletion of UBR5 HECT domain

Early B cell development occurs in the BM compartment so we examined BM B cells in 6-week-old *Mb1*^*WT/CRE*^;*Ubr5*^*ΔHECT/∆HECT*^ mice. In *Ubr5*^*ΔHECT/ΔHECT*^ and *Ubr5*^*ΔHECT/WT*^ *mice*, the number of total BM cells and frequency of B220^+^ B cells showed no significant difference compared to their WT littermates (Figure 2A-C; Supplemental Figure 1A-B). Further analysis of specific subtypes of B cells revealed a striking decrease in IgD^+^ mature B cell populations (Figure 2D-E; Supplemental Figure 1C-D). Decreases in mature B cells were compensated in BM by a slight increase in propre B cell population (Figure 2F-G). Further breakdown of the population revealed an increase in pro B, but not pre B cell population (Figure 2F-I). These studies demonstrate an impact to mature cells within BM, as well as changes to composition of pro and pre B cells following loss of HECT domain.

**Figure 2.**
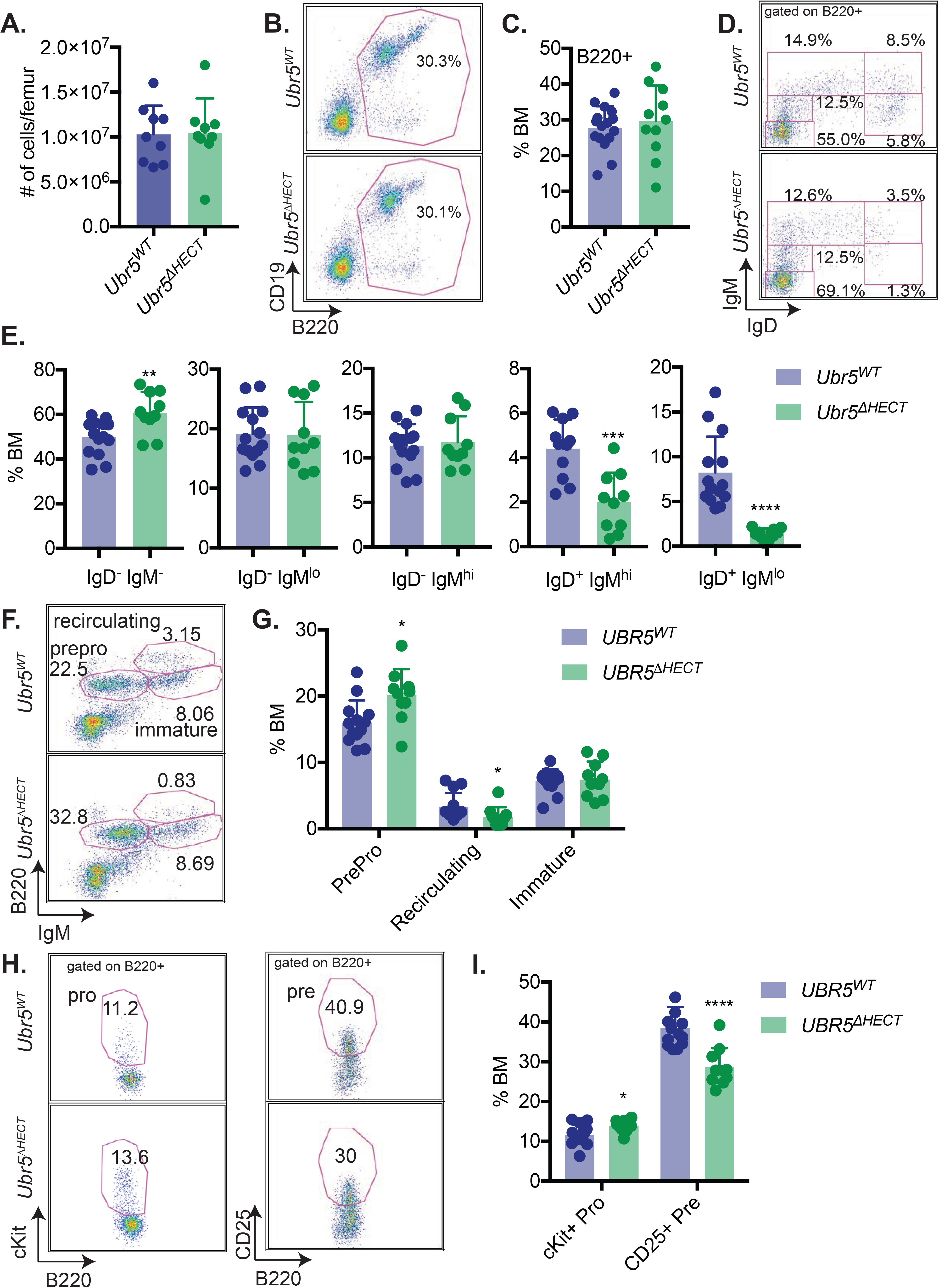
The loss of the HECT domain of *Ubr5* leads to decreased numbers of mature B cells within the BM. (A) Bar graph of the total number of cells per femur. (B) Representative flowcytometry plot of total B220^+^ in the BM. (C) Bar graph of the frequency of B220^+^ cells per femur. (D) Representative flowcytometry plots gated on B220^+^ cells for pro and pre B cells (B220^+^IgM^−^ IgD^−^), immature B cells (B220^+^IgM^lo^IgD^−^), transitional B cells (B220^+^IgM^+^IgD^−^), early mature B cells (B220^+^IgM^+^IgD^+^), and late mature B cells (B220^+^IgM^−^IgD^+^). (E) Bar graphs of B cell populations shown in D. (F) Representative flow cytometry plots gated for pro and pre B cells (B220^+^IgM^−^), immature B cells (B220^+^IgM^lo^), and recirculating B cells (B220^+^IgM^+^). (G) Bar graph of the population breakdown shown in F. (H) A representative flowcytometry plot gated on B220^+^ cells gating for pro B cells (B220^+^IgM^−^c-kit^+^) and pre B cells (B220^+^IgM^−^CD25^+^) (I) Bar graph representing the population breakdown shown in H. (N=10, * P<0.05, ** P<0.01, *** P<0.001)

Following development of B cells in BM, cells migrate to the spleen where they undergo maturation and activation.^23^ *Mb1^CRE/WT^;Ubr5^ΔHECT/ΔHECT^* mice have smaller spleens as well as a reduction in number of total splenocytes (Figure 3A-B). Although the mice had a total reduction of splenocytes, frequency of B220^+^ cells in both WT littermates and *Ubr5*^*ΔHECT/ΔHECT*^ mice was ~45% (Figure 3C), and splenic architecture was unaltered following loss of the HECT domain of *Ubr5* (Figure 3D). In addition, the transitional B cell population stages T1, T2, and T3 in the spleen had no significant differences in *Ubr5*^*ΔHECT/ΔHECT*^ mice versus WT littermates (Figure 3 E-F). However, there was significant impact on mature B1 and B2 subsets within the spleen. We found that in the B2 subset, marginal zone B cells were significantly reduced from approximately 10% of B220^+^ splenocytes to 2% in *Ubr5*^*ΔHECT/ΔHECT*^ mice whereas the follicular B cell compartment frequency was slightly increased, despite a reduction in absolute number of follicular B cells in *Ubr5*^*ΔHECT/ΔHECT*^ mice (Figure 3H-I; Supplemental Figure 1E-F).

**Figure 3.**
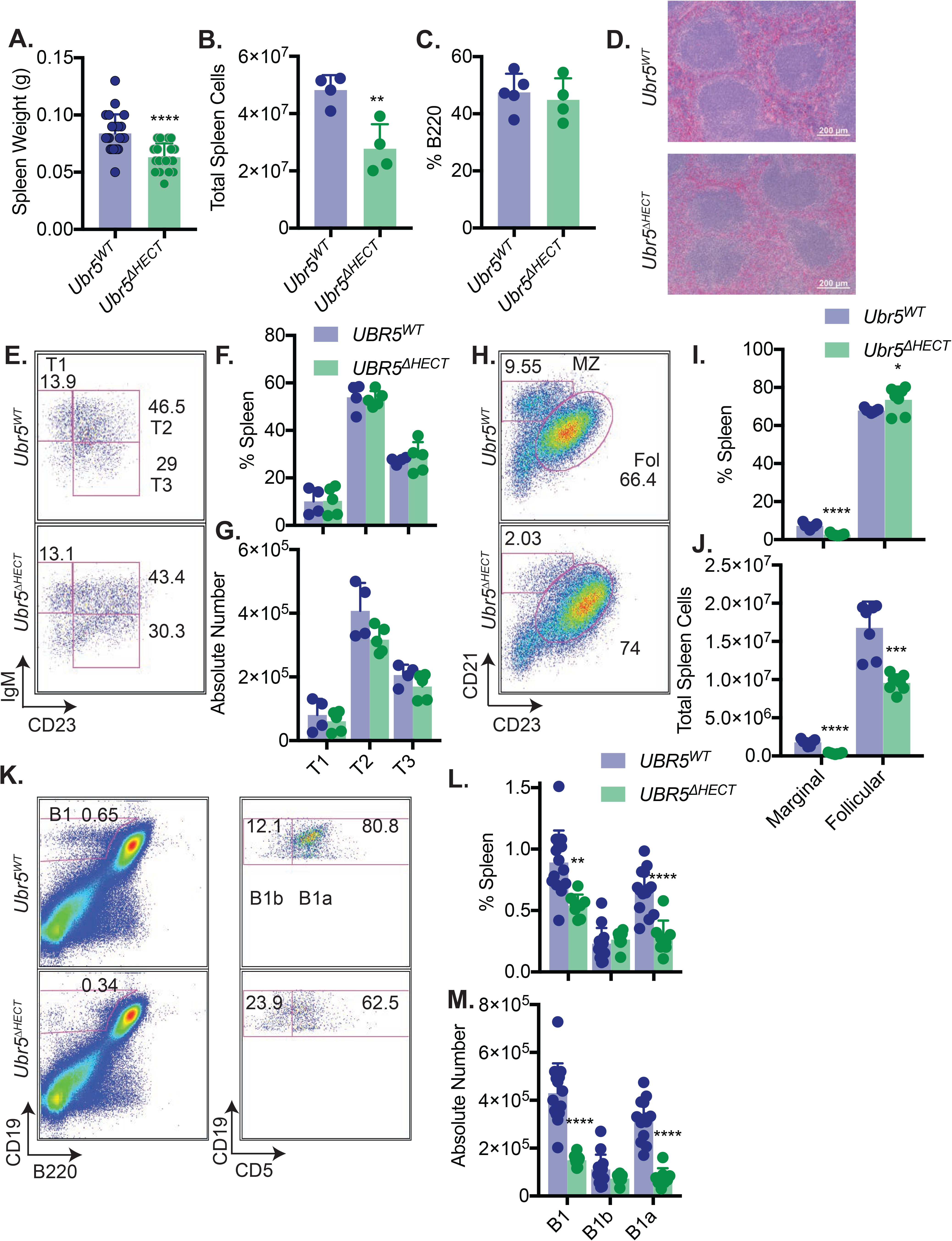
The loss of the HECT domain of *Ubr5* leads to reduction in marginal and B1a splenic cells, but no effects on transitional B cells. (A) Bar graph of spleen weight. (N=19) (B) Bar graph of the total number of splenocytes. (N=4) (C) Bar graph of frequency of B220^+^ cells per spleen. (N=4) (D) An example hematoxylin and eosin stain of spleens. (N=4) (E) Representative flow cytometry plot gated on B220^+^ for transitional cells: T1 cells (B220^+^CD93^+^IgM^+^CD23^−^), T2 cells (B220^+^CD93^+^IgM^+^CD23^+^), and T3 cells (B220^+^CD93^+^IgM^−^CD23^+^). (N=5) (F-G) Bar graphs of percentages and absolute number of transitional cells within the spleen. (H) Representative flowcytometry plots gated on B220^+^ for follicular B cells (B220^+^CD21^+^CD23^+^) and marginal zone B cells (B220^+^CD21^+^CD23^−^). (N=7) (I-J) Bar graphs of percentages and absolute number of follicular and marginal zone B cells within the spleen. (K) Representative flowcytometry plots gated for B1a (B220^+^CD19^lo^CD5^+^) and B1b cells (B220^+^CD19^lo^CD5^−^). (N=8) (L & M) Bar graph of percentages and absolute number of B1a and B1b cells. (* P<0.05, ** P<0.01, *** P<0.001, **** P<0.0001)

The B1 population is responsible for innate immunity and is the first line of defense for infection. In spleen, the B1 population was reduced by almost 2-fold in *Ubr5*^ΔHECT/ΔHECT^ mice and when further separated into B1a and B1b, the reduction was exclusively found in the B1a subpopulation (Figure 3K-M). We further evaluated B1 populations in the peritoneal cavity and found a ~75% reduction in B1 cells, mainly from a reduction of B1a cells (Supplemental Figure 2). These findings demonstrate a significant loss of populations required for innate immunity with loss of B1 and marginal zone B cells following deletion of the HECT domain of *Ubr5*.

### Alterations in follicular B cell subsets and activation of B cells

Evaluating cell surface markers on the follicular B cell population, we found that *Ubr5*^*ΔHECT/ΔHECT*^ follicular B cells had abnormal protein expression with low IgD and high IgM compared to their WT littermates (Figure 4A-B). The follicular population in *Ubr5*^*ΔHECT/ΔHECT*^ mice also had high CD23 protein expression, but normal expression of CD5 and CD1d on follicular B cells (Figure 4A-B). To further define alterations in the B cell compartment, we analyzed cell cycle status. While follicular B cells are typically in the resting state, *Ubr5*^*ΔHECT/ΔHECT*^ cells are more quiescent and had increased cells in G0 for both transitional and mature B cells (Supplemental Figure 3A). Additionally, staining with cellular proliferation marker Ki67 on spleen sections showed a reduction of Ki67 staining specifically in white pulp of spleens from *Ubr5*^*ΔHECT/ΔHECT*^ mice (Supplemental Figure 3B). These studies demonstrated alterations in mature spleen cells with both phenotypic changes and cell cycle alterations.

**Figure 4.**
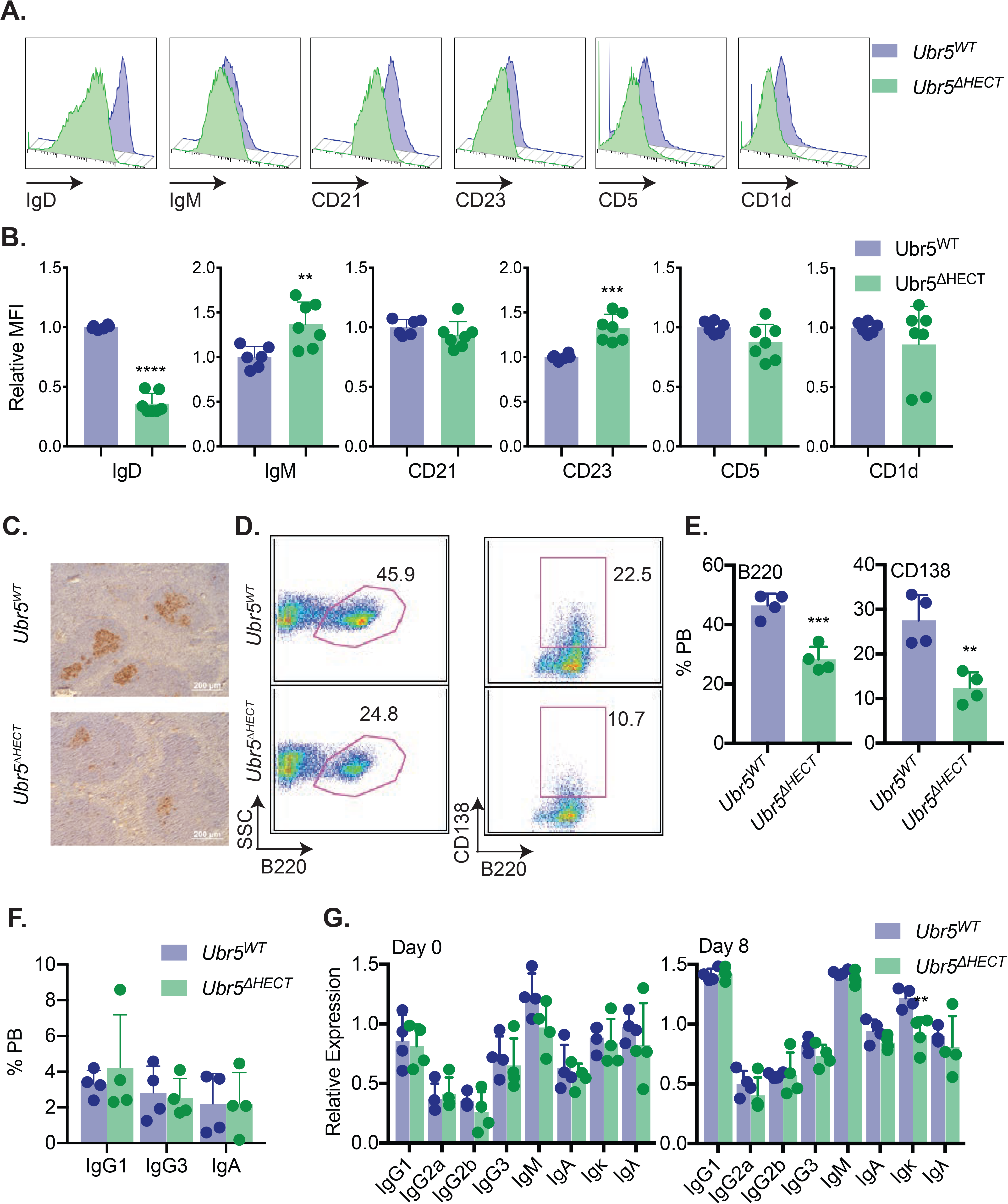
The follicular B cells in *Ubr5^∆HECT^* mice have altered phenotype and diminished differentiation capabilities. (A) Representative histogram of cell surface markers of *Ubr5^WT^* and *Ubr5^∆HECT^* follicular B cells (B220^+^CD21^+^CD23^+^). (B) Bar graphs of relative MFI of follicular B cells depicted in A. (C) Representative PNA IHC staining of spleen 7 days post stimulation with SRBC. (D) Representative flowcytometry plots of B220^+^ and CD138^+^ plasma cells in the peripheral blood. (E) Bar graphs representing percentages of B220^+^ and CD138^+^ cells in the peripheral blood depicted in D. (F) Bar graphs representing percentages of IgG1^+^, IgG3^+^, and IgA^+^ B220^+^ cells in the peripheral blood. (G) Bar graphs of ELISA indicating relative expression for different immunoglobulin types before (Day 0) and after (Day 8) immune system stimulation with SRBC. (N=4) (* P<0.05, ** P<0.01, *** P<0.001, **** P<0.0001)

Activation of follicular B cells by T dependent antigens leads to formation of germinal centers (GC) in secondary lymphoid tissues and generation of antibody producing plasma cells. In order to determine if *Ubr5*^*ΔHECT/ΔHECT*^ follicular B cells have normal function and whether germinal center formation can be induced, we immunized 6-week-old mice with SRBC. Eight days following immunization, we performed immunohistochemical (IHC) analysis on spleens with peanut agglutinin, a GC marker. Staining revealed a significant decrease in overall number and size of GCs in *Ubr5^∆HECT/∆HECT^* mice, but in white pulp with GCs, there were multiple small GCs (Figure 4C). Only ~65% of white pulp in spleens of *Ubr5*^*ΔHECT/ΔHECT*^ mice contained GCs compared to ~83% in WT littermate controls. We evaluated the ability of follicular B cells to terminally differentiate into plasma cells within PB and found a ~50% reduction of B cells and plasma cells (Figure 4D-E; Supplemental Figure 1G-H). Although we had a significant decrease in CD138^+^ antibody producing cells, levels of IgG1, IgG3, and IgA were not changed in unstimulated *Ubr5^∆HECT/∆HECT^* mice (Figure 4F), and measurement of antibodies IgG, IgM, and IgA within sera were unaltered suggesting that *Ubr5* HECT domain deletion does not affect basal antibody levels (Figure 4G). These studies indicate an important role of Ubr5 in the function and activation of mature B cells.

### Loss of UBR5 HECT domain leads to increased expression of spliceosome components found to interact with UBR5

The HECT domain of UBR5 is thought to be required for its ubiquitination activity implicating deletion could lead to increased accumulation of UBR5 substrates targeted for degradation by the proteasome. To quantitatively determine protein differences in *Ubr5*^*ΔHECT/ΔHECT*^ versus WT littermates, we labeled splenic B220^+^ cells with tandem mass tags (TMT), combined samples in equal concentrations, and analyzed by MS (Figure 5A). Proteomic analysis identified 15,584 unique peptides, 2,797 quantifiable proteins, and 1,675 proteins with greater than or equal to three unique peptides. We identified 143 proteins that were either significantly decreased or enriched in *Ubr5*^*ΔHECT/ΔHECT*^ B220^+^ splenocytes (Figure 5B and Supplementary Table 2). Principle component analysis (PCA) plot showed protein isolated from 3 independent mice from either genotype clustered together, but that the two independent genotypes clustered separately (Figure 5C). In addition, identified proteins were distributed in different cellular compartments (Figure 5D). Of the differentially expressed proteins, 104 ≥1.3 fold significantly overexpressed proteins were enriched for proteins associated with mRNA processing, RNA splicing, and mRNA splicing via the spliceosome, whereas the 39 ≤0.70 fold significantly decreased proteins in spleen B220^+^ cells from *Ubr5*^ΔHECT/ΔHECT^ were proteins associated with immune system process and protein transport (Figure 5E-G). As suggested by flowcytometry, IgD protein was the highest down-regulated protein following deletion of *Ubr5* HECT domain (Figure 5E). CD22, which is associated with B cell activation, was also down-regulated in correlation with the follicular B cell phenotype in *Ubr5*^*ΔHECT/ΔHECT*^ mice (Figure 5E). Intriguingly, the second highest expressed protein in splenic B cells of *Ubr5*^*ΔHECT/ΔHECT*^ was UBR5 (Figure 5E-F). Proteomic profiling revealed an increase in proteins associated with mRNA splicing in B cells lacking HECT domain of UBR5 suggesting a novel role of UBR5 in mRNA splicing.

**Figure 5.**
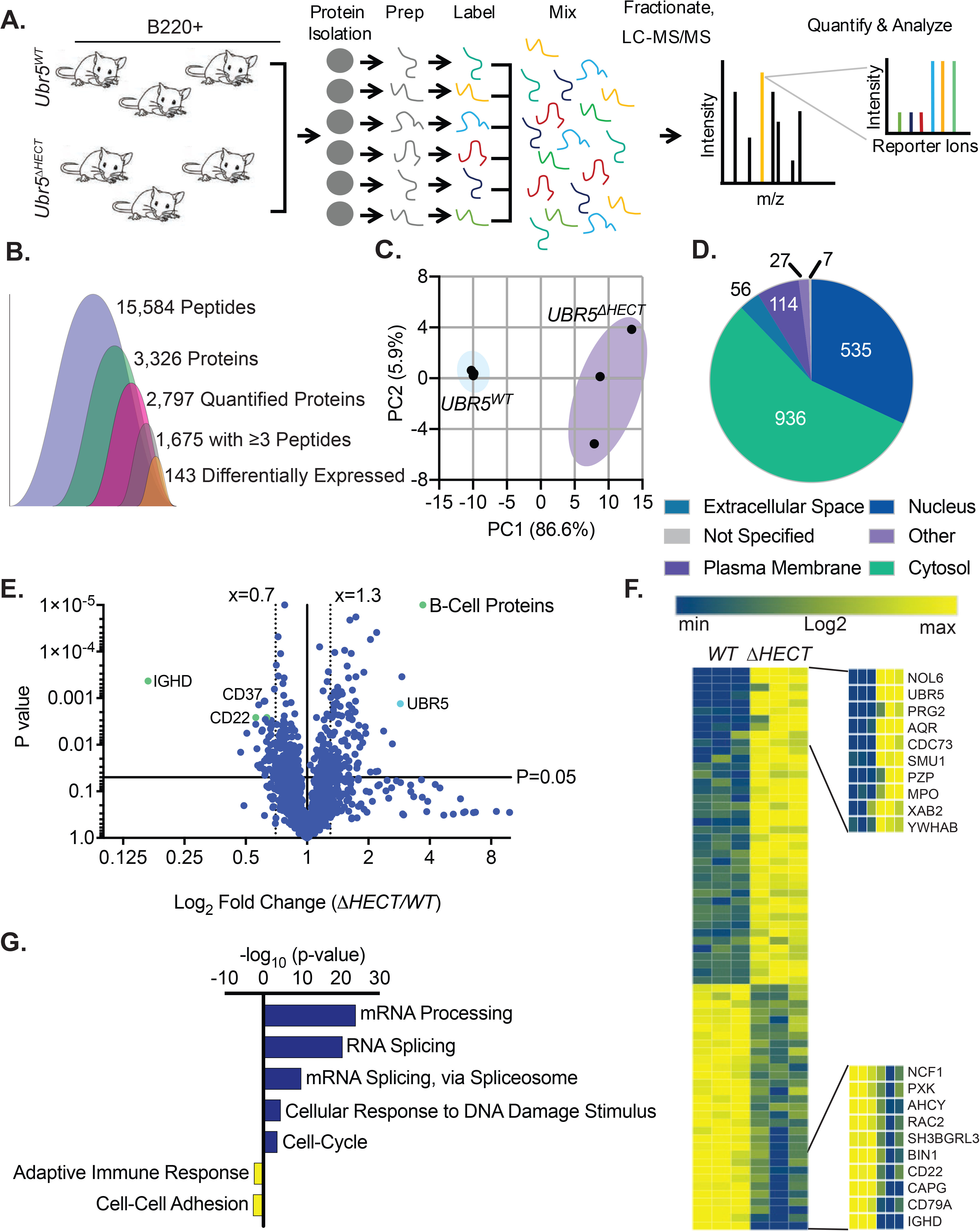
Loss of HECT domain results in a reduction in B cell development proteins and an enrichment in proteins regulating mRNA splicing. (A) Schematic of sample preparation of *UBR5^WT^*and *Ubr5^∆HECT^* spleens for TMT MS analysis. (N=3). (B) Quantification of identified peptides and proteins. (C) PCA using components 1 and 2 showing clustering of *WT* and ΔHECT samples for significantly up-regulated proteins. (D) Pie chart of localizations for all 2,786 proteins identified by mass spec. (E) Volcano plot of the Log_2_ fold change *WT/ΔHECT* showing proteins significantly up-regulated (p<0.05) 1.3-fold or more and significantly (p<0.05) down-regulated 0.7-fold or more. (F) Heatmap of the significantly up-regulated and significantly down-regulated proteins showing the top 10 up-regulated and top 10 down-regulated proteins. (G) Gene ontology analysis showing pathways known to be associated with significantly up-regulated (≥1.3, p≤0.05) and significantly down-regulated (≤0.7, p≤0.05) proteins using DAVID 6.8 software. For up-regulated proteins, includes pathways with ≥10 proteins associated and a p-value ≤0.01. Down-regulated proteins pathways with p-value ≤0.05.

To identify putative substrates and interacting partners of UBR5 in MCL, we performed immunoprecipitation followed by mass spectrometry (IP-MS) utilizing human MCL patient derived cell lines, Jeko1 and Mino (Figure 6A).^24, 25^ We identified 115 proteins with ≥3 unique peptides and, similar to our TMT analysis, gene ontology analysis showed proteins enriched in mRNA splicing via the spliceosome (Figure 6B and Supplementary Table 3). Comparing MS datasets from TMT labeled proteins and endogenous IP-MS revealed 89 proteins overlapping and seven proteins were up-regulated in the TMT study including UBR5 (Figure 7 D-E). Intriguingly, all six of the other identified proteins are associated with mRNA splicing (SF3B3, SMC2, PRPF8, DHX15, SNRNP200, and EFTUD2). These proteins are classified as core spliceosome components including U2 (SF3B3) and associated U5 small nuclear ribonucleoprotein (snRNP) complex (EFTUD2, SNRNP200, and PRPF8) (Figure 6E). Of the identified proteins, none have previously been characterized as UBR5 interactors or substrates, suggesting a novel pathway of UBR5 regulation and the identification of potential novel UBR5 substrates.

**Figure 6.**
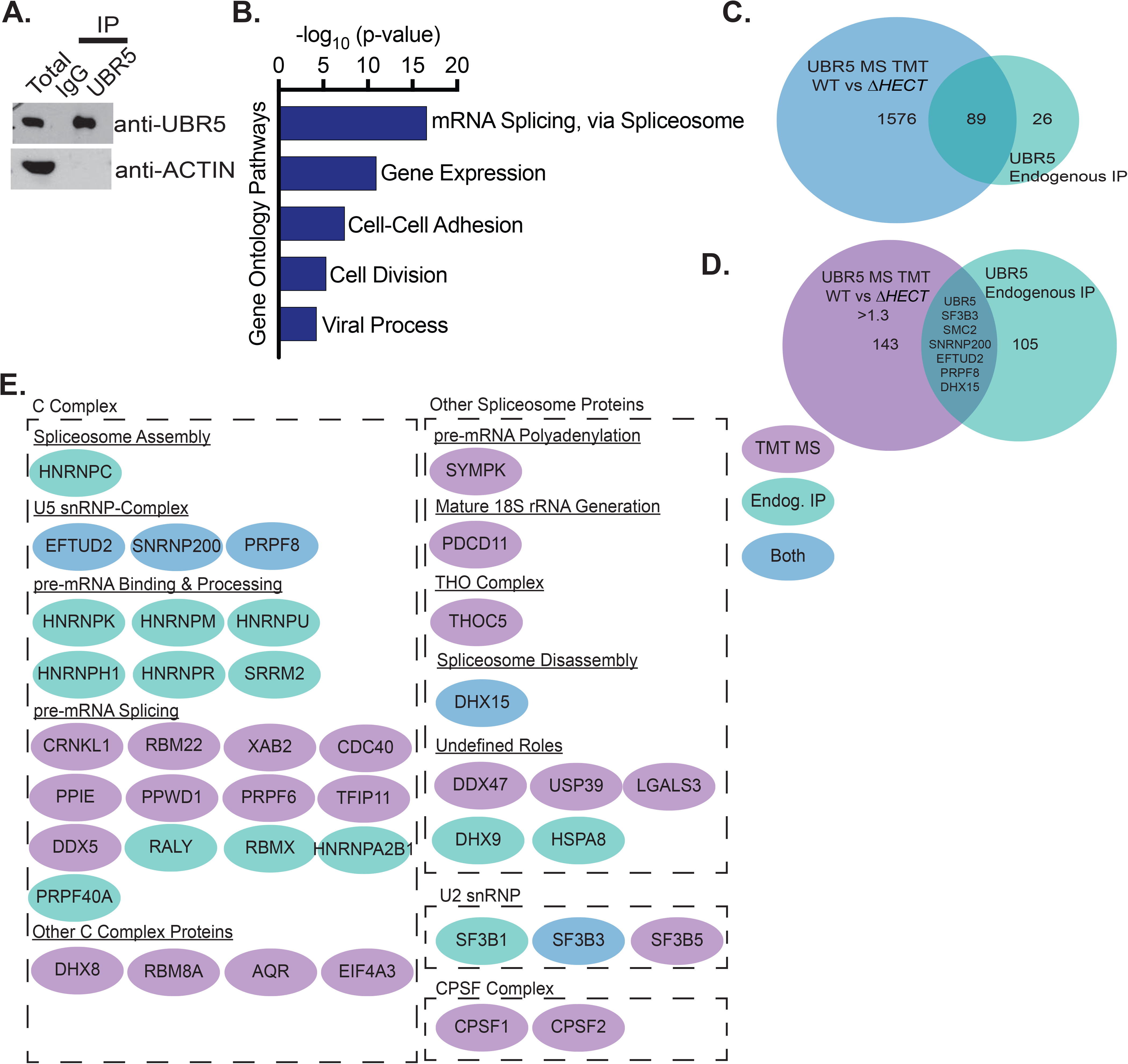
UBR5 interacts with spliceosome components. (A) Western blot of immunoprecipitation of UBR5 from JEKO1 MCL cell line used for MS analysis. MS experiment was performed in duplicate (B) Gene ontology analysis of immunoprecipitated proteins. (C) Venn diagram showing the significant overlap of proteins identified in the TMT labelled *UBR5^WT^* and *Ubr5^ΔHECT^* MS and those identified in the UBR5 endogenous IP MS. (D) Venn diagram showing overlap of the proteins significantly up-regulated in the *Ubr5^∆HECT^* samples and those identified in the UBR5 endogenous IP. (E) Spliceosome associated proteins found in endogenous immunoprecipitation and/or up-regulated in spleen B220^+^ cells.

**Figure 7.**
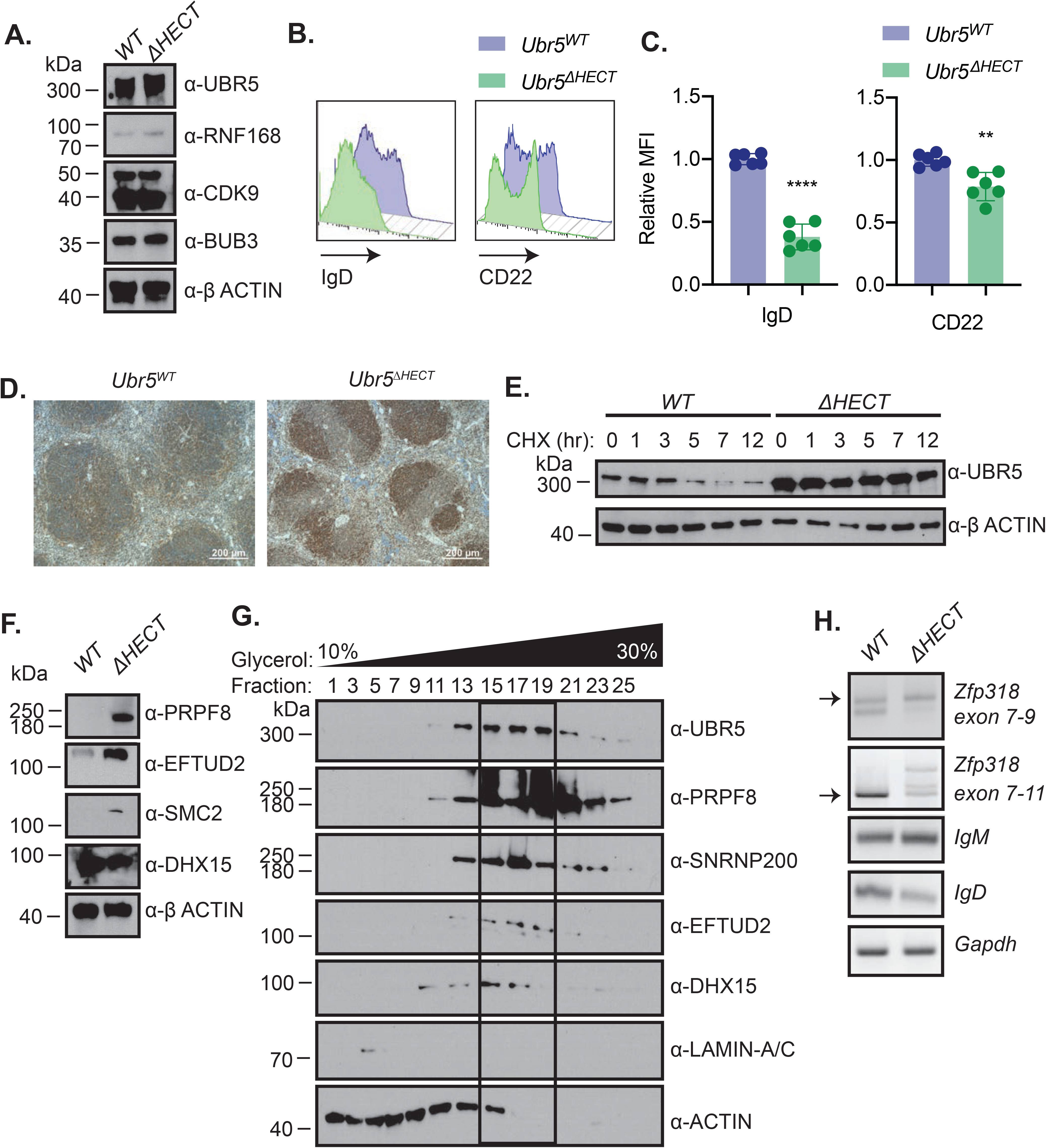
Loss of HECT domain leads to aberrant splicing. (A) Western blot of known UBR5 substrates in spleen B cells (B220^+^) *Ubr5^W^*^T^ and *Ubr5^∆HECT^* mice. (B) Representative histogram of cell surface markers of spleen cells (B220^+^). (C) Bar graphs of relative MFI of the markers in B. (D) Representative immunohistochemistry staining of UBR5 protein in spleen. (E) Western blot of UBR5 in follicular B cells (B220^+^CD21^+^CD23^+^) *Ubr5^W^*^T^ and *Ubr5^∆HECT^* after treatment with 10µg/mL cycloheximide. (F) Western blot of proteins identified in MS in spleenic B cells (B220^+^) *Ubr5^WT^* versus *Ubr5^∆HECT^* mice. (G) 10-30% glycerol fractionation of nuclear lysate from HEK293T cells followed by western blot for UBR5 and spliceosome components. (H) RT-PCR of mRNA in *Ubr5^WT^* and *Ubr5^∆HECT^* follicular B cells (B220^+^CD21^+^CD23^+^). Arrows point to the bands of interest. (N=3) (** P<0.01, **** P<0.0001).

### Alterations in splicing factors and splicing in B cells

UBR5 has been implicated in a number of cellular processes including cell cycle (CDK9), DNA damage (RNF168), and cell division (BUB3).^5-7, 26^ BUB3 was found in the quantitative MS analysis; however, accumulation was not seen in cells with deletion of *Ubr5* HECT domain (Supplementary Table 1). In addition, western blot confirmed that BUB3, CDK9, and RNF-168 did not accumulate in *Ubr5*^*ΔHECT/ΔHECT*^ B220^+^ splenocytes (Figure 7A). Flowcytometry validated cell surface markers associated with activation and maturation, CD22 and IgD, expression was significantly decreased in total B220^+^ splenocytes (Figure 7B-C). UBR5 was the second highest expressed protein in the MS; however, none of the identified peptides had coverage within the HECT domain (data not shown). To confirm overexpression of UBR5 protein, we performed IHC and western blot analysis and found higher protein expression of UBR5 in *Ubr5*^*ΔHECT/ΔHECT*^ spleens (Figure 7A and D). In order to determine if loss of HECT domain leads to stabilization of the protein, we performed half-life analysis with cycloheximide treatment. *Ubr5*^*ΔHECT/ΔHECT*^ B220^+^ splenocytes had higher protein expression due to stabilization of the protein (Figure 7E). These studies validate the MS findings that known substrates do not accumulate and loss of the HECT domain leads to overexpression and stabilization of UBR5.

To evaluate protein overexpression of RNA splicing components, we performed western blot analysis on B220^+^ splenocytes. *Ubr5*^*ΔHECT/ΔHECT*^ B220^+^ splenocytes had increased expression of EFTUD2, SMC2, and PRPF8 (Figure 7F). Because three of the identified proteins EFTUD2, SNRNP200, and PRPF8 are part of the U5 snRNP complex, we wanted to determine whether UBR5 elutes with the U5 complex. We performed glycerol density gradient and found that EFTUD2, SNRNP200, PRPF8, and UBR5 eluted in fractions 15-19 suggesting they are found in the same complex (Figure 7G).

Expression of *IgM* and *IgD* is regulated by alternative splicing of the Ig heavy chain locus. As shown in the follicular B cell population, *IgM* and *IgD* have aberrant expression suggesting defects in splicing. To confirm altered splicing, we isolated follicular B cells from *Ubr5*^*ΔHECT/ΔHECT*^ and WT littermates for analysis of transcript levels. PCR confirmed increased *IgM* transcripts and decreased *IgD* transcripts in *Ubr5*^*ΔHECT/ΔHECT*^ follicular B cells (Figure 7H). It was previously shown that mRNA of Zinc Finger Protein 318 (*Zfp318*) has two known splice variants.^27^ It is thought that the longer transcript form containing exon 1-10 plays a key role in alternative splicing of immunoglobulin heavy chain locus. We found decreased expression of transcript spliced from exon 7 to exon 11 and aberrant variant transcripts in *Ubr5*^*ΔHECT/ΔHECT*^ follicular B cells, and decreased transcript containing exon 7 to exon 9 suggesting expression of *Zfp318* splice variants may also be contributing to B cell defects (Figure 7H). These studies demonstrate that loss of HECT domain leads to increased expression of spliceosome components and aberrant splicing.

## DISCUSSION

In this report, we demonstrate that ubiquitin E3 ligase UBR5 plays a key role in B cell maturation and activation via regulation of alternative splicing. While UBR5 is present in all B cell populations, deletion of *Ubr5* HECT domain had limited identifiable impact on development of BM B cell populations. This suggests a crucial role for UBR5 in maturation of B cells. With the loss of the UBR5 HECT domain, there is normal frequency of transitional B cells that migrate to the spleen; although there is a block in differentiation of naïve B cells which corresponds to the MCL tumor population. The greatest impact is on B1a and marginal zone B cells that are significantly reduced within the spleen and peritoneal cavity suggesting impaired innate immunity. Although follicular B cell population frequency is only slightly impacted, these cells are phenotypically abnormal and have a reduced capacity to generate plasma cells.

Interrogation of global proteomic changes in the splenic B cell population demonstrates a novel role of UBR5 in mRNA splicing via the spliceosome. The spliceosome plays an important role in providing genetic diversity. In immune cells, alternative splicing is suggested to play a key role in B cell differentiation, activation, and survival.^28^ A number of important pathways and genes in the immune system require alternative splicing. In human B cells, ~90% of genes with multiple exons undergo alternative splicing.^29^ The spliceosome involves multicomponent complexes including five snRNPs that coordinate mRNA splicing.^30^ Our finding that components of the spliceosome are both up-regulated and found interacting with UBR5 in MCL cell lines identifies a new pathway of UBR5 involvement. The striking phenotype of IgD in HECT domain mutants supports defects in splicing since IgM and IgD are generated from alternative splicing of the heavy chain gene.^31^ It is well established that IgM and IgD play a key role in immunity and are required for B cell receptor signaling and immune response, but little is known regarding how alternative splicing of *IgD* and *IgM* occur. In addition, markers of mature naïve B cells, *CD21* and *CD23*, have multiple splice variants regulated by alternative splicing.^32, 33^ Our studies find that in transitional and follicular B cell populations, *CD23* has altered expression, which may be due to splicing. Importantly, expression of SF3B3 identified in our MS has been suggested to lead to alternative splicing of *Ezh2* and promote tumorigenesis.^34^ In addition, EZH2 has been previously shown to play a key role in germinal center formation and EZH2 mutations promote lymphoid transformation.^35^ Furthermore, SF3B1, another component of the U2 snRNP complex that interacts with SF3B3 is frequently mutated in chronic lymphoblastic leukemia and a variant of Myelodysplastic Syndrome, suggesting that dysregulation of this complex may be important for development of hematopoietic malignancies.^36, 37^ Although these studies suggest a role of UBR5 in the spliceosome, further investigation is required to determine if ubiquitination activity is required or whether UBR5 acts as a scaffolding protein during splicing.

Loss of the HECT domain of UBR5 leads to stabilization and overexpression which could be due to UBR5 potentially undergoing self-ubiquitination.^38^ Also, UBR5 is predicted to bind RNA further supporting its role in RNA splicing.^38^ More importantly these studies identifying the consequences of UBR5 HECT domain loss on alternative splicing in the immune system not only reveal mechanisms regulating B cells development, but since ~18% of MCL patient genomes contain UBR5 mutations, it may lead to further understanding of lymphoma transformation and provide a potential therapeutic target. Overall, our findings reveal a novel mechanism of regulation by UBR5 via alternative mRNA splicing, provides evidence that the ubiquitin E3 ligase is a novel regulator of B cell maturation, and suggests a role of UBR5 in lymphoma transformation.

## AUTHOR CONTRIBUTIONS

S.S., T.G, and S.M.B. conceived and designed the experiments. S.S., T.G., H.V., J.H.G, M.R.G. and S.M.B preformed experiments and analysis. S.S., T.G., R.W.H., and S.M.B. wrote the manuscript. H.C.H.L. and N.T.W. provided technical and material support. All authors reviewed the manuscript before submission.

## ACKNOWLEDGEMENTS

We would like to thank Dr. M. Reth for the use of the *Mb1*^*CRE*^ mice; Dr. S. Korolov for helpful discussion and providing *Mb1*^*CRE*^ mice; and Patrick Swanson, Creighton University for helpful discussions. Also, we would like to thank the UNMC FlowCytometry Research Facility, UNMC Mouse Genome Engineering Core Facility, and UNMC Mass Spectrometry and Proteomics Core Facility for expert assistance. The core facilities are administrated through the Office of the Vice Chancellor for Research and supported by state funds from the Nebraska Research Initiative (NRI) and The Fred and Pamela Buffett Cancer Center’s National Cancer Institute Cancer Support Grant. S.M.B. and N.T.W. are supported by the National Institutes of Health (P20GM121316) and ACS institutional grant. This study was in part supported by the Frances E. Lageschulte & Evelyn B. Weese New Frontiers Medical Research Fund. W.H.S. is supported by the UNMC NIH training grant (5T32CA009476-23). This publication was supported by the Fred & Pamela Buffett Cancer Center Support Grant from the National Cancer Institute under award number P30 CA036727.

