## Supplemental Material for "E3 Ligase UBR5 HECT domain mutations in lymphoma control maturation of B cells via alternative splicing"

### Swenson et al Supplemental Figure 1

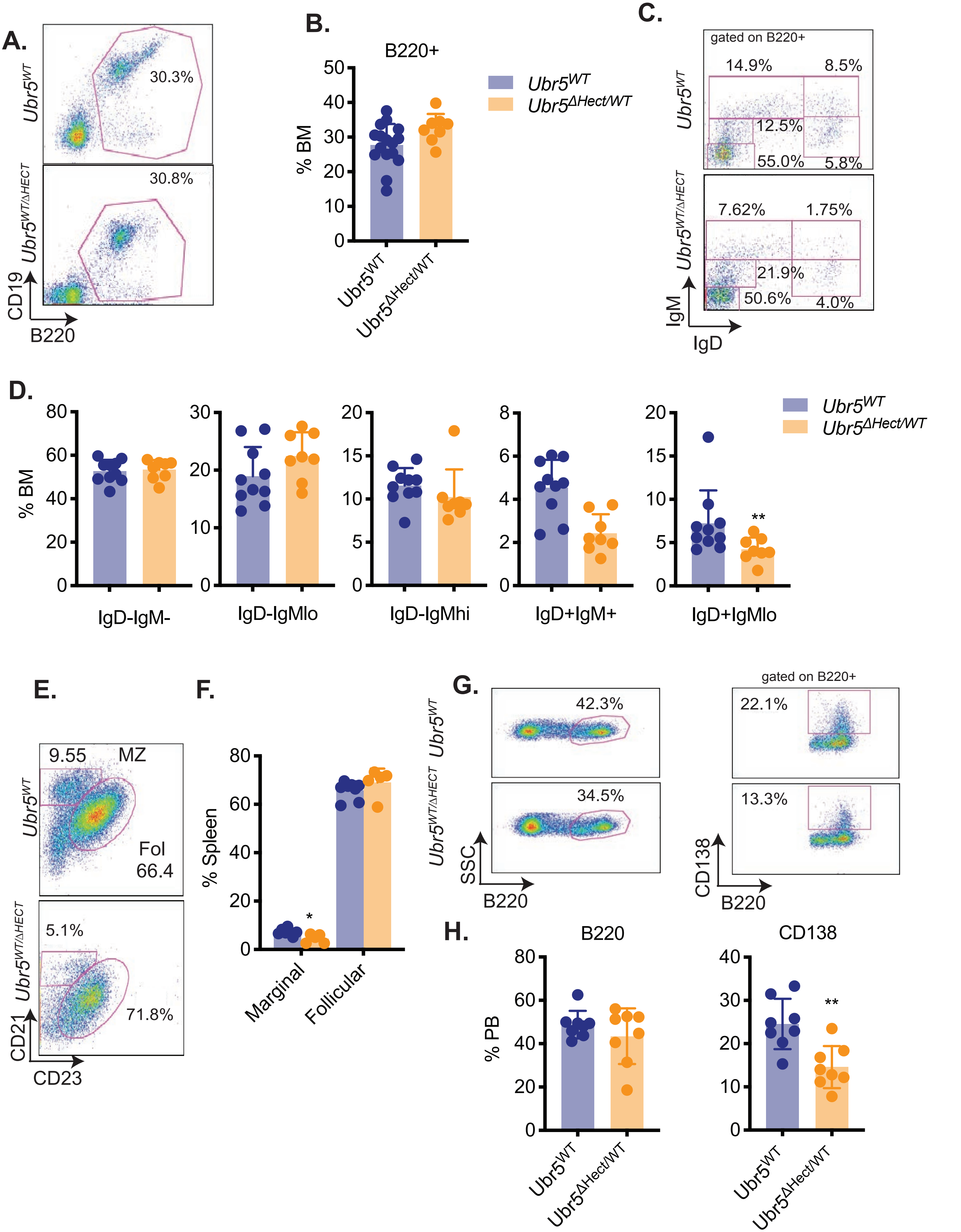

### Swenson et al Supplemental Figure 2

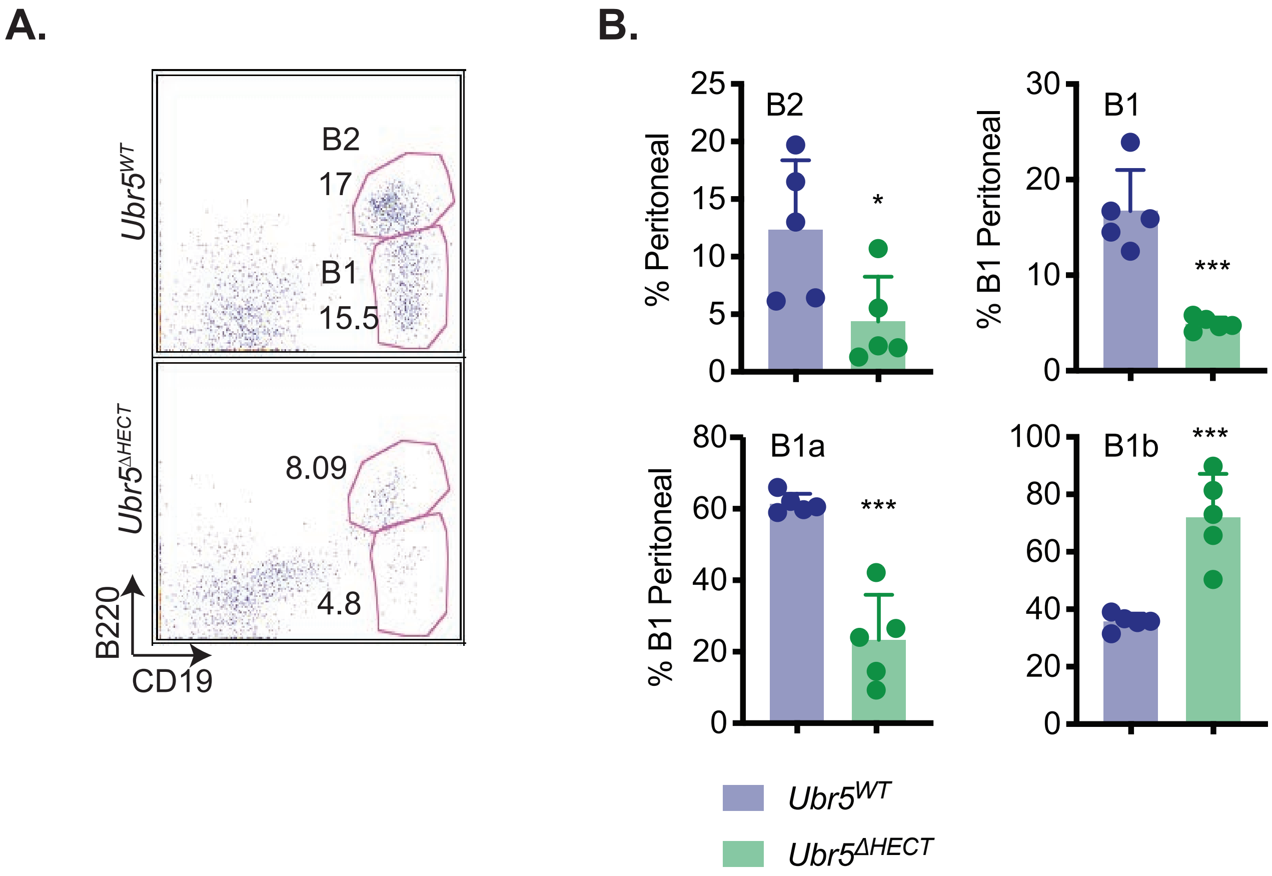

### Swenson et al Supplemental Figure 3

A.

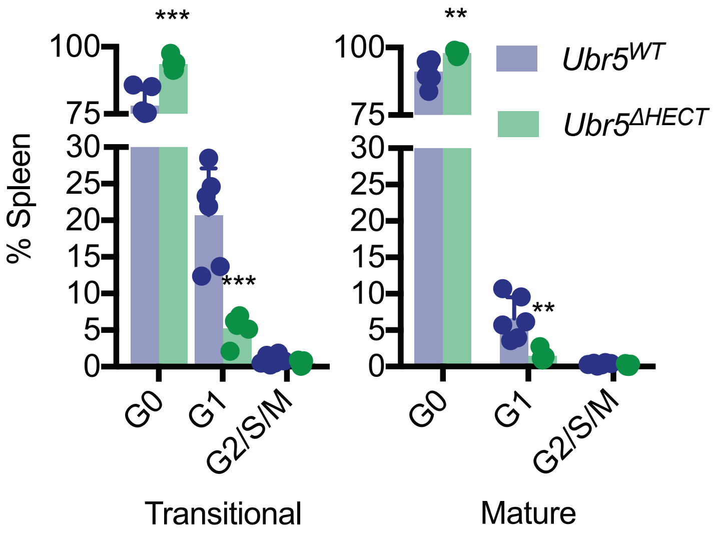


B.

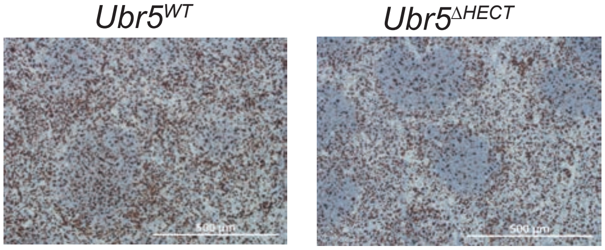

**Swenson et al. Supplemental Data**

#### Supplemental Figure Legends

**Supplemental Figure1. *Ubr5*$^{\Delta HECT/WT}$ mice do not have a pronounced phenotype compared to *Ubr5*$^{WT}$ mice.** (A) Representative flowcytometry plot of total B220$^{+}$ in the BM. (B) Bar graph of the frequency of B220$^{+}$ per femur (N=8). (C) Representative flowcytometry plots gated on B220$^{+}$ cells for pro and pre B cells (B220$^{+}$IgM$^{-}$IgD$^{-}$), immature B cells (B220$^{+}$IgM$^{lo}$IgD$^{-}$), transitional B cells (B220$^{+}$IgM$^{+}$IgD$^{-}$), early mature B cells ((B220$^{+}$IgM$^{+}$IgD$^{+}$), and late mature B cells (B220$^{+}$IgM$^{-}$IgD$^{+}$). (D) Bar graphs of B cell populations shown in C (N=8). (E) Representative flowcytometry plots gated on B220$^{+}$ for follicular B cells (B220$^{+}$CD21$^{+}$CD23$^{+}$) and marginal zone B cells (B220$^{+}$CD21$^{+}$CD23$^{-}$). (F) Bar graph of the percentage of follicular and marginal zone B cells within the spleen (N=8). (G) Representative flowcytometry plots of B220$^{+}$ and CD138$^{+}$ in the peripheral blood. (H) Bar graphs representing percentages of B220$^{+}$ and CD138$^{+}$ in the peripheral blood depicted in G (N=8, * P<0.05, ** P<0.01, *** P<0.001).

**Supplemental Figure 2. Loss of the Ubr5 HECT domain leads to decreased B1a percentage and increased B1b percentage within the peritoneal cavity.** (A) Representative FACS plot of total B220$^{+}$ cells in the peritoneal cavity. (B) Bar graphs representing the percentages of B2 (B220$^{+}$CD19$^{+}$), B1 (B220$^{lo}$CD19$^{+}$), B1b (B220$^{+}$CD19$^{lo}$CD5$^{-}$), and B1a (B220$^{+}$CD19$^{lo}$CD5$^{+}$) (N=5, * P<0.05, ** P<0.01, *** P<0.001).

**Supplemental Figure 3. Loss of HECT domain leads to increased splenic cells in G0.**

(A) Cell cycle breakdown of transitional ($B220^+IgM^+IgD^-$) and mature ($B220^+IgM^+IgD^+$) splenocytes. (B) Ki67 IHC staining of spleen (N=3, * P<0.05, ** P<0.01, *** P<0.001).

#### Supplemental Material and Methods

| Genotyping Primer | Sequence |
|---|---|
| Ubr5 3' F | 5' – GCT CCT CCA GTT CAA ACG GT – 3' |
| Ubr5 3' R | 5' – GAC CCA AAC AAT GAC TCA GAA C – 3' |
| Ubr5 5' F | 5' – GGA TTA AAG GTG TGC GCT GCC – 3' |
| Mb1 F | 5' – CCC GCC TCA CCT GTG AAA A – 3' |
| Mb1 R | 5' – CTG GCA CCA CTG CAC AGA A – 3' |
| Pan Cre F | 5' – CCC AAG AAG AAG AGG AAA GTC – 3' |
| Pan Cre R | 5' – AGA CCA GGC CAG GTA TCT C – 3' |

| qRT-PCR Primer | Sequence |
|---|---|
| Gapdh F | 5' – CAT GGC CTT CCG TGT TCC TA – 3' |
| Gapdh R | 5' – CTG GTC CTC AGT GTA GCC CAA – 3' |
| Ubr5 F | 5' – TGA GGT TTC TAC GAT CTG TGG C – 3' |
| Ubr5 R | 5' – AAA CAC ACG TTT GCA TTT TCC A – 3' |
| Ubr5 Hect F | 5' – CTC CAG TTC AAA CGG TGG TT – 3' |
| Ubr5 Hect R | 5' – CTG GCA ATG ATG GGC TAG AT – 3' |

IHC Antibodies:

| Antibody | Company | Ref# |
|---|---|---|
| UBR5 | Abcam | ab70311 |
| PNA | Vector Laboratories | B-1075 |

Western Blot Antibodies:

| Antibody | Company | Ref# |
|---|---|---|
| UBR5 | Cell Signaling | 65344S |
| BUB3 | Abcam | ab133699 |
| RNF168 | Santa Cruz | sc-101125 |
| CDK9 | Cell Signaling | 2316S |
| EFTUD2 | Bethyl | A300-957A |
| SNRNP200 | Abcam | ab176715 |

| | | |
|---|---|---|
| ACTIN | Santa Cruz | sc-47778 |
| DHX15 | Protein Tech | 12265-1-AP |
| PRPF8 | Abcam | ab79237 |
| SMC2 | Cell Signaling | 5394S |
| LAMIN-A/C | Bethyl | A303-431A |

Flow Cytometry Antibodies:

| Antibody | Fluorochrome | Clone | Company |
|---|---|---|---|
| B220 | BV421, BV510, APC Cy7 | RA3-6B2 | BioLegend |
| CD19 | APC Cy7, PE Cy7 | 6D5 | BioLegend |
| IgM | PE, PE Cy7, FITC | RMM-1 | BioLegend |
| IgD | APC, BV711 | 11-26c.2a | BioLegend |
| cKit | APC | 2B8 | BioLegend |
| CD25 | FITC | 3C7 | BioLegend |
| CD21/CD35 | PE | 7E9 | BioLegend |
| CD23 | APC, Alexa 700 | B3B4, B3B4 | BioLegend |
| CD5 | FITC, Per cp Cy 5.5 | 53-7.3 | BioLegend |
| CD1d | Pac Blue | 1B1 | BioLegend |
| CD138 | BV421 | 281-2 | BioLegend |
| CD22 | APC | OX-97 | BioLegend |
| CD93 | FITC | AA4.1 | BioLegend |
| DAPI | | | |
| Ki67 | FITC | 16A8 | BioLegend |

Mass Spectrometry Method:

Samples were loaded onto trap column Acclaim PepMap 100 75µm x 2 cm C18 LC Columns (Thermo Scientific™) at flow rate of 5µl/min then separated with a Thermo RSLC Ultimate 3000 (Thermo Scientific™) from 5-20% solvent B (0.1% FA in 80% ACN) from 10-98 minutes at 300nL/min and 50ºC with a 120 minutes total run time for fractions one and two. For fractions three to six, solvent B was used at 5-45% for the same duration. Eluted peptides were analyzed by a Thermo Orbitrap Fusion Lumos Tribrid (Thermo Scientific™) mass spectrometer in a data dependent acquisition mode using synchronous precursor selection method. A survey full scan MS (from m/z 375-

1500) was acquired in the Orbitrap with a resolution of 120,000. The AGC target for MS2 in iontrap was set as $1x10^4$ and ion filling time set as 150ms and fragmented using CID fragmentation with 35% normalized collision energy. The AGC target for MS3 in orbitrap was set as $1x10^5$ and ion filling time set as 200ms with a scan range of 100-500 and fragmented using HCD with 65% normalized collision energy. Protein identification was performed using proteome discoverer software version 2.2 (Thermo Fisher Scientific) by searching MS/MS data against the UniProt mouse protein database. The search was set up for full tryptic peptides with a maximum of 2 missed cleavage sites. Oxidation, TMT6plex of the amino terminus, GG and GGQ ubiquitination, phosphorylation, and acetylation were included as variable modifications and carbamidomethylation and TMT6plex of the amino terminus were set as fixed modifications. The precursor mass tolerance threshold was set at 10ppm for a maximum fragment mass error of 0.6 Da with a minimum peptide length of 6 and a maximum peptide length of 144. The significance threshold of the ion score was calculated based on a false discovery rate calculated using the percolator node. Protein accessions were put into Ingenuity Pathway Analysis (QIAGEN Inc.) to identify gene symbols and localizations. Gene ontology pathway analysis was performed using DAVID Bioinformatics Database 6.8 using the functional annotation tool.
